# Protective effect and molecular mechanisms of human non-neutralizing cross-reactive spike antibodies elicited by SARS-CoV-2 mRNA vaccination

**DOI:** 10.1101/2024.02.28.582613

**Authors:** Jordan Clark, Irene Hoxie, Daniel C. Adelsberg, Iden A. Sapse, Robert Andreata-Santos, Jeremy S. Yong, Fatima Amanat, Johnstone Tcheou, Ariel Raskin, Gagandeep Singh, Irene González-Domínguez, Julia E. Edgar, Stylianos Bournazos, Weina Sun, Juan Manuel Carreño, Viviana Simon, Ali H. Ellebedy, Goran Bajic, Florian Krammer

**Affiliations:** Department of Microbiology, Icahn School of Medicine at Mount Sinai, New York, NY, USA; Center for Vaccine Research and Pandemic Preparedness (C-VARPP), Icahn School of Medicine at Mount Sinai, New York, NY, USA; Retrovirology Laboratory, Department of Microbiology, Immunology and Parasitology, Paulista School of Medicine, Federal University of São Paulo (UNIFESP), São Paulo, SP, Brazil; Laboratory of Molecular Genetics and Immunology, The Rockefeller University, New York, NY, USA; Department of Pathology, Molecular and Cell Based Medicine, Icahn School of Medicine at Mount Sinai, New York, NY, USA; Division of Infectious Diseases, Department of Medicine, Icahn School of Medicine at Mount Sinai, New York, NY, USA; The Global Health and Emerging Pathogens Institute, Icahn School of Medicine at Mount Sinai, New York, NY, USA; Department of Pathology and Immunology, Washington University School of Medicine, St. Louis, MO 63110, USA; Center for Vaccines and Immunity to Microbial Pathogens, Washington University School of Medicine, St. Louis, MO 63110, USA; The Andrew M. and Jane M. Bursky Center for Human Immunology and Immunotherapy Programs, Washington University School of Medicine, Saint Louis, MO 63110, USA

## Abstract

Neutralizing antibodies correlate with protection against SARS-CoV-2. Recent studies, however, show that binding antibody titers, in the absence of robust neutralizing activity, also correlate with protection from disease progression. Non-neutralizing antibodies cannot directly protect from infection but may recruit effector cells thus contribute to the clearance of infected cells. Also, they often bind conserved epitopes across multiple variants. We characterized 42 human mAbs from COVID-19 vaccinated individuals. Most of these antibodies exhibited no neutralizing activity *in vitro* but several non-neutralizing antibodies protected against lethal challenge with SARS-CoV-2 in different animal models. A subset of those mAbs showed a clear dependence on Fc-mediated effector functions. We determined the structures of three non-neutralizing antibodies with two targeting the RBD, and one that targeting the SD1 region. Our data confirms the real-world observation in humans that non-neutralizing antibodies to SARS-CoV-2 can be protective.

## Introduction

Severe acute respiratory syndrome coronavirus 2 (SARS-CoV-2) emerged in Wuhan, China in December 2019 and is the causative agent of the coronavirus disease 2019 (COVID-19) pandemic. As of November 2023, there have been over 6.9 million official deaths worldwide due to COVID-19 but the current estimate based on excess mortality is as high as 27.3 million^1^. In late 2020, two mRNA-based vaccines, BNT162b2 (Pfizer) and mRNA-1273 (Moderna) received emergency use authorization from the United States Food and Drug Administration (FDA) and by several countries worldwide in the following months. Numerous additional vaccines based on different vaccine platforms are now available, including adenovirus-based vectors, recombinant protein vaccines, DNA vaccines, and inactivated vaccines^2^. Most of these vaccines utilize the spike glycoprotein of SARS-CoV-2 as an antigen, which is present on virions as a trimer. Spike protomers are synthesized as single polypeptide chains which are subsequently cleaved into two subunits, S1 and S2, by host proteases such as furin^3^. The receptor binding domain (RBD) within S1 interacts with the angiotensin-converting enzyme 2 (ACE2) receptor, which is used by the virus to enter host cells^4^. Antibodies elicited against the spike, particularly against the RBD, can therefore prevent viral attachment and neutralize the virus. However, neutralizing epitopes have also been reported within the spike protein’s S2 and N-terminal domains (NTD)^5–10^. While it has been shown that neutralizing titers correlate with protection from COVID-19^11–14^, the protective roles played by other non-neutralizing antibodies are not well understood. Several reports have highlighted that vaccination elicits a larger number of non-neutralizing antibodies than infection^15^, and that despite the well-documented loss of neutralizing activity against recently arisen variants of concern (VOCs), individuals vaccinated with the BNT162b2 and mRNA-1273 vaccines still exhibit protection from disease^16^. Of note, Earle and colleagues found that binding antibody titers alone correlate with protection^14^, and a more recent study showed that binding antibodies correlated with protection from progression to severe disease, even when neutralizing titers are significantly reduced^17^. Taken together, these data suggest that other factors aside from neutralization are important for the control and clearance of infection. Whilst non-neutralizing antibodies are unable to prevent SARS-CoV-2 entry by binding virions via their fragment antibody binding (Fab) regions, their fragment crystallizable (Fc) regions can still be engaged by the Fc gamma receptors (FcγRs) present on various immune effector cell populations^18^. Interactions with these receptors can mediate protection via antibody-dependent cellular cytotoxicity (ADCC)^19^ and antibody-dependent phagocytosis (ADCP)^20^, as well as via modulation of antiviral CD8 T cell responses. Furthermore, the Fc regions can be recognized by the complement component C1q^21^. This binding initiates the complement cascade and forms the membrane attack complex (MAC), which can lyse infected cells and destroy virions^21^. ADCC is typically mediated by natural killer (NK) cells^22^, although monocytes and macrophages have also been implicated in this process^23^, whereas ADCP is mediated by monocytes, macrophages, neutrophils, and eosinophils^20^. In a previous study, we identified 42 SARS-CoV-2 spike binding monoclonal antibodies (mAbs) derived from plasmablasts of three adults who had completed their primary BNT162b2 vaccination^15^. Of these, 12 bound RBD, 10 bound NTD, 17 bound S2, and three bound unknown epitopes. Interestingly, only seven mAbs (16%) were neutralizing. Several studies have highlighted the importance of effector cell-mediated immunity in protecting against SARS-CoV-2 infection^24–29^. Therefore, here we investigated whether these non-neutralizing antibodies could contribute to protection from SARS-CoV-2 infection in a lethal animal model and determined cryo-EM structures of selected mAbs to understand their molecular mechanism of action.

## Results

### Non-neutralizing antibodies maintain binding to divergent spike proteins

We previously cloned 42 monoclonal antibodies from the plasmablasts of three subjects who had received two doses of the Pfizer mRNA vaccine^10^. Due to the rapid evolution of immune evasive SARS-CoV-2 variants of concern (VOCs), the binding of these mAbs was tested against the spike proteins of the ancestral WA-1, Alpha (B.1.1.7), Beta (B.1.351), Delta (B.1.617.2), and the Omicron subvariants BA.5 and XBB.1. Many of the antibodies retained binding to the SARS-CoV-2 variants (Fig. 1A). Of the seven neutralizing mAbs tested, four NTD-binding mAbs exhibited reduced binding to the VOC spikes. Of these, PVI.V6-2 exhibited abrogated binding to Beta, BA.5, and XBB.1 spike, while PVI.V6-14^30^ lost binding to BA.5 and XBB.1 spike. PVI.V5-6 and PVI.V6-7 maintained spike binding, however, the minimum binding concentration increased against all the tested VOCs. Of the two RBD-binding, neutralizing mAbs, PVI.V6-4 displayed reduced binding to BA.5 and XBB.1 spike, while PVI.V3-9 retained binding to each of the VOCs. Similarly, the NTD binding mAb PVI.6-11 bound strongly to all the VOC spikes. Conversely, binding was preserved for the majority of the non-neutralizing mAbs, with only four mAbs demonstrating a reduction in spike binding. To investigate whether the neutralizing antibodies that maintained binding to the VOCs could still neutralize, microneutralization assays were carried out using authentic ancestral, B.1.351, B.1.1617.2, BA.1, and BA.5 viruses. Regardless of spike binding, neutralization of the mAbs was abolished against all tested VOCs, except in the case of PVI.V6-4 and PVI.V3-9, which retained neutralization against Delta (Fig. 1B).

**Figure 1.**
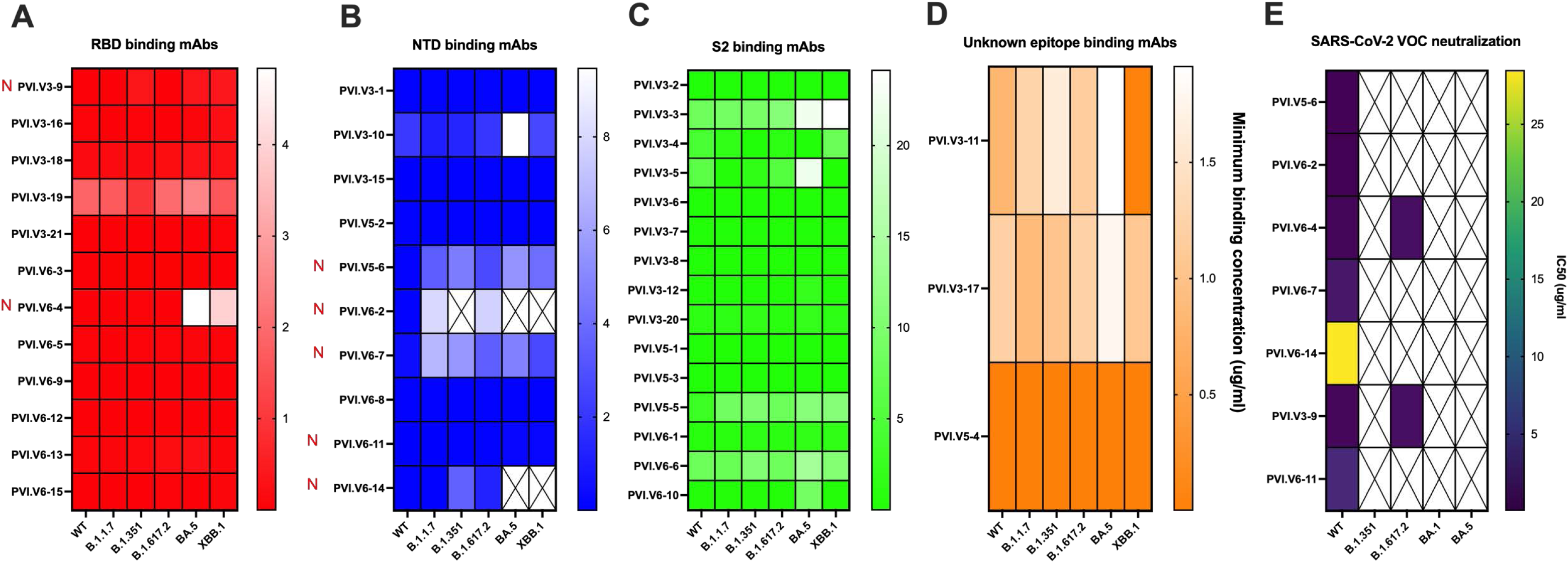
Characterization of mAb binding and neutralization against SARS-CoV-2 variants. **(A-D)** Binding of RBD-specific **(A)**, NTD-specific **(B)**, S2-specific **(C)**, and unknown epitope-specific **(D)** SARS-CoV-2 mAbs to the spike proteins of SARS-CoV-2 variants. Neutralizing mAbs are denoted with red Ns. **(E)** Neutralizing activity of the neutralizing mAbs against authentic SARS-CoV-2 variants.

### Non-neutralizing antibodies protect from lethal challenge murine challenge and reduce lung viral titers

To determine the protective abilities of the mAbs against SARS-CoV-2 infection, BALB/cAnNHsd mice were treated intraperitoneally with 10mg/kg of antibody and infected intranasally with 5x the 50% lethal dose (LD_50_) of mouse-adapted SARS-CoV-2 (MA10)^31^. Of the RBD binding mAbs, the two neutralizing antibodies were found to confer 100% survival, wherein animals treated with PVI.V6-4 lost no weight and animals treated with PVI.V3-9 lost approximately 5% body weight (Fig. 2A). Three non-neutralizing mAbs (PVI.V3-2, PVI.V6-3, and PVI.V6-12) were also found to confer 100% survival while two others (PVI.V3-16 and PVI.V6-9) conferred 80% survival. Of the mAbs binding NTD, most of the neutralizing mAbs were found to promote 100% survival with less than 5% weight loss (PVI.V-5-6, PVI.V6-2, PVI.V6-7, and PVI.V6-11) with one mAb (PVI.V6-14) eliciting 60% protection with around 12% weight loss (Fig. 2B). One anti-NTD non-neutralizing antibody (PVI.V5-2) promoted 100% survival with approximately 15% weight loss. Of the mAbs cloned from the three mRNA recipients, the majority were found to be non-neutralizing antibodies targeting S2. Of these, PVI.V5-1 conferred 100% survival with no weight loss, while three others (PVI.V3-4, PVI.V3-13, and PVI.V6-6) elicited 80% survival; treatment with one mAb, PVI.V3-8, resulted in 60% survival (Fig. 2C). Another antibody that binds a so far unknown epitope outside of the RBD, NTD, and S2 (PVI.V5-4), was also found to promote 100% survival following lethal challenge (Fig. 2D). In total, 19 of the 42 mAbs (45%) displayed protection from disease, with 12/19 (63%) of these being non-neutralizing mAbs. To determine whether the protective non-neutralizing mAbs may be weakly neutralizing, microneutralization assays were repeated using an increased mAb concentration of 200 µg/ml. These assays identified mAb PVI.V6-3 as a weak neutralizer, with an IC50 of 98.86 µg/ml (Fig. S1). As the murine challenge experiment was carried out using a dose of 10 mg/kg, the neutralizing effects of this mAb at the *in vivo* available concentration are likely negligible.

**Figure 2.**
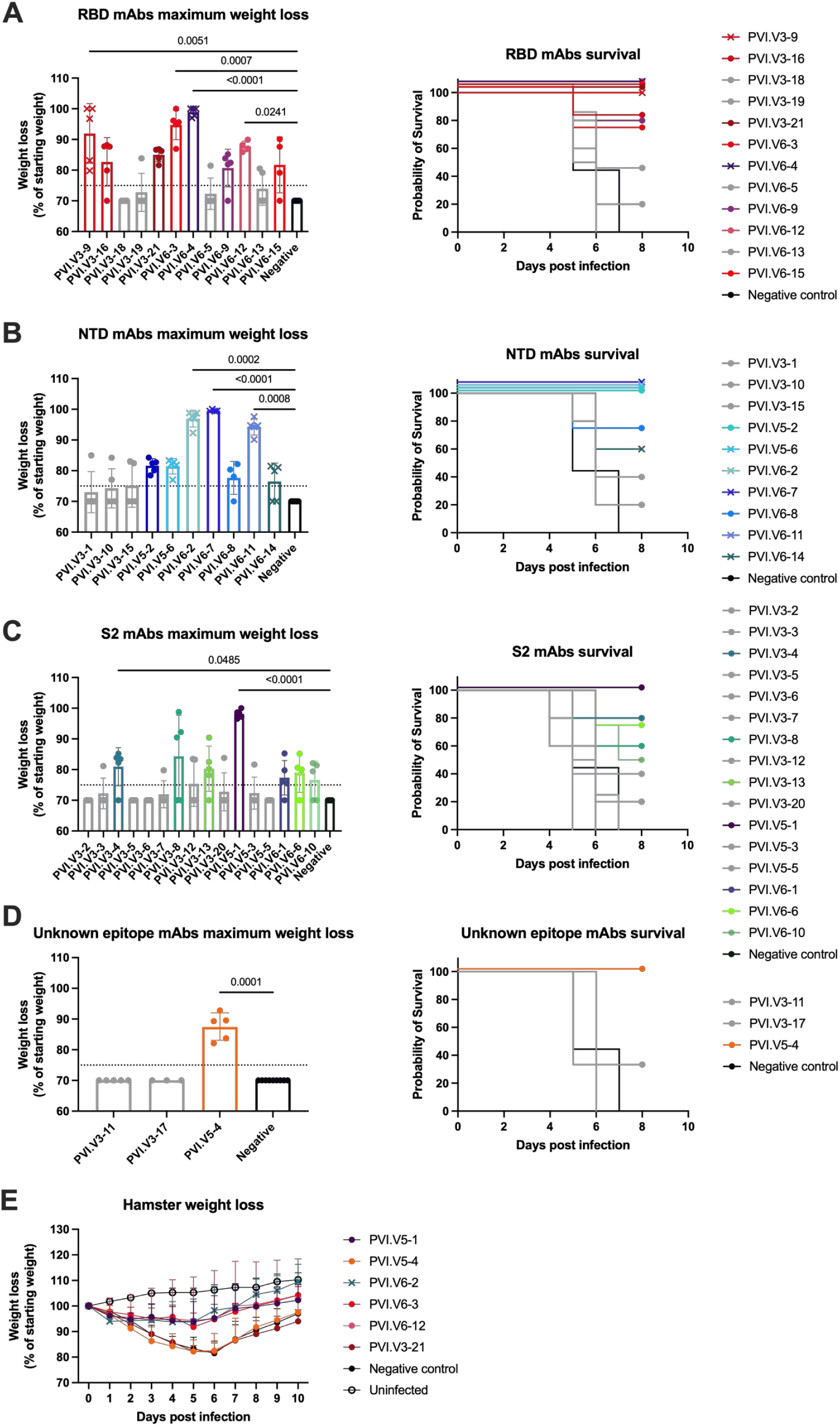
Investigating the protection elicited by the mAbs using lethal animal challenge models. **(A-D)** Maximum weight loss and survival of BALB/cAnNHsd mice treated with RBD-specific **(A)**, NTD-specific **(B)**, S2-specific **(C)**, and unknown epitope-specific **(D)** SARS-CoV-2 mAbs prior to lethal challenge with mouse-adapted SARS-CoV-2. **(E)** Hamster weight loss following treatment with the 6 most protective mAbs and challenge with wild type SARS-CoV-2. Neutralizing mAbs are shown as crosses, while non-neutralizing mAbs are shown as circles. In **(A-D)**, the vertical bars in the maximum weight loss graphs indicates the mean with standard deviation. P values are shown only for statistically significant comparisons as determined by a Kruskal-Wallis with Dunn’s multiple comparisons test.

To test these mAbs further using a severe model of SARS-CoV-2 infection, the transgenic hACE2-K18 mouse model was utilized with ancestral SARS-CoV-2. This mouse model expresses human ACE2, rendering the mice highly susceptible to SARS-CoV-2 infection which progresses to fatal neurotropic infection. We therefore tested whether the seven most protective mAbs (PVI.V5-1, PVI.V5-4, PVI.V6-2, PVI.V6-3, PVI.V6-12, and PVI.V3-21) were able to confer protection when administered 2 hours prior to infection with 3x and 5xLD_50_ doses of ancestral SARS-CoV-2. At a 3xLD_50_ infectious dose, the neutralizing mAb PVI.V6-2, and the non-neutralizing mAb PVI.V5-1 promoted 100% survival, while PVI.V6-3 and PVI.V6-12 promoted 80% and 60% survival respectively (Fig. S2). The non-neutralizing mAbs PVI.V5-2, PVI.V5-4, PVI.V3-21 conferred 40% survival. At the 5xLD_50_ dose only PVI.V6-2 and PVI.V6-3 protected the mice from challenge, with mice treated with the other non-neutralizing mAbs succumbing to infection by 8-10 dpi (Fig. S2). We then sought to validate the protective effects of these mAbs using a different model which more closely recapitulates human disease. To this end, we tested whether PVI.V5-1, PVI.V5-4, PVI.V6-2, PVI.V6-3, PVI.V6-12, and PVI.V3-21 were able to protect against ancestral SARS-CoV-2 challenge in the Golden Syrian hamster model. Hamsters were treated intraperitoneally with 10mg/kg mAb 2 hours prior to intranasal infection with wild-type SARS-CoV-2. Treatment with all mAbs, except for PVI.V5-4 and PVI.V3-21, resulted in decreased weight loss compared to an irrelevant negative control mAb, indicating that these mAbs can protect against wild-type SARS-CoV-2 infection in an alternative animal model with wild-type virus (Fig. 2E).

To determine whether the 19 protective mAbs could reduce viral titers, mice were treated with 10mg/kg of antibody, infected using a 1xLD_50_ dose of MA10 to prevent mortality, and lungs were harvested at 3- and 5-days post-infection (dpi) for viral enumeration via plaque assay. At 3dpi all mAbs except for PVI.V5-1, PVI.V5-2, PVI.V6-1, and PVI.V5-6 were found to significantly reduce lung viral titers (Fig. 3A-D). Mice treated with the neutralizing antibodies PVI.V6-4, PVI.V3-9, and PVI.V6-11 exhibited no detectable virus in their lungs whereas other neutralizing antibodies targeting NTD only moderately reduced viral titers. The RBD, S2, and unknown epitope-binding, non-neutralizing mAbs were found to reduce viral titers ranging from 10- to 1000-fold. At 5dpi all of the mAbs were found to significantly reduce viral titers with the majority lowering titers below the limit of detection, indicating that both the neutralizing and non-neutralizing mAbs promote viral clearance (Fig. 3E-H).

**Figure 3.**
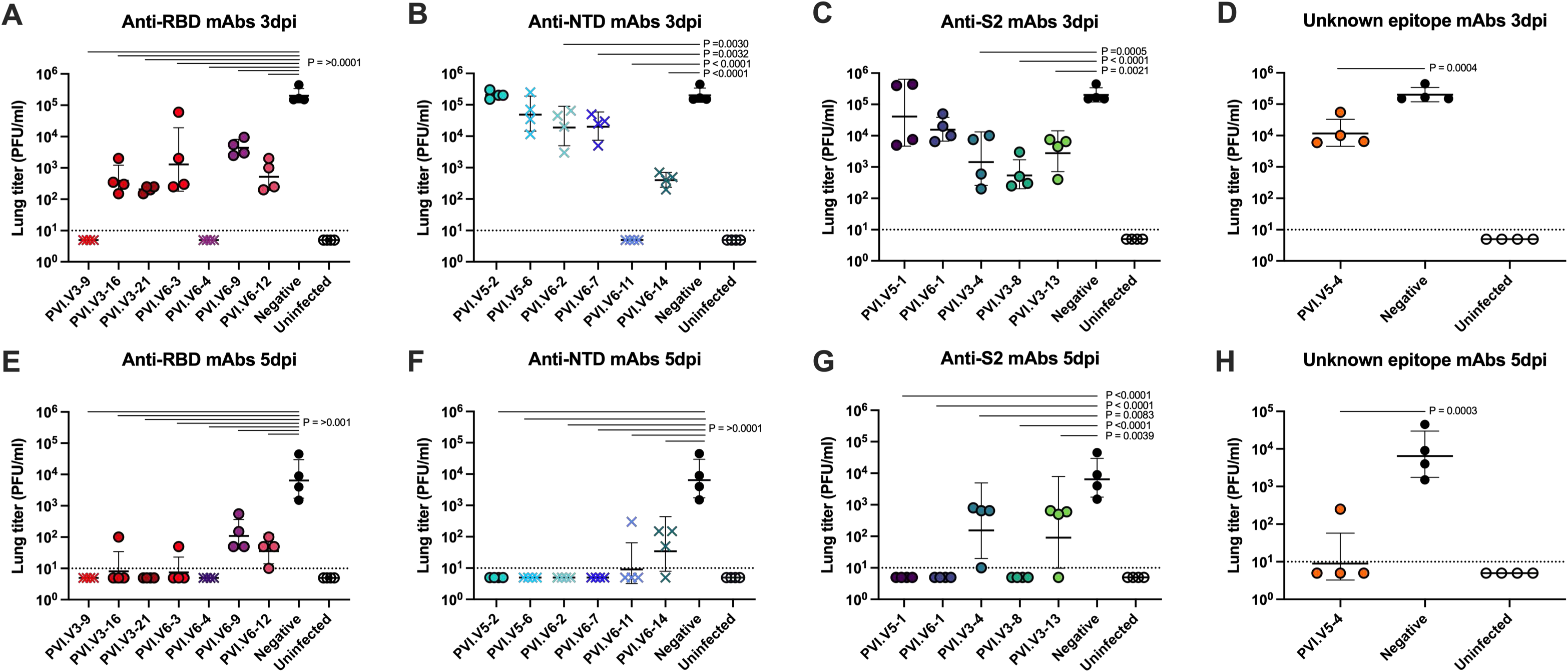
Mouse lung titers following mAb treatment and SARS-CoV-2 challenge. **(A-D)** murine lung titers 3 dpi after treatment with RBD-specific **(A)**, NTD-specific **(B)**, S2-specific **(C)**, and unknown epitope-specific **(D)** SARS-CoV-2 mAbs and challenge with mouse adapted SARS-CoV-2. **(E-H)** lung titers 5 dpi after treatment with RBD-specific **(E)**, NTD-specific **(F)**, S2-specific **(G)**, and unknown epitope-specific **(H)** SARS-CoV-2 mAbs and challenge with mouse adapted SARS-CoV-2. Neutralizing mAbs are shown as crosses, while non-neutralizing mAbs are shown as circles. P values are shown only for groups with titers statistically significant compared to the negative control, as determined by an ordinary one-way ANOVA with Dunnett’s multiple comparisons test with a single pooled variance.

### Structural epitope mapping of non-neutralizing epitopes

Given the protective potency of the non-neutralizing mAbs PVI.V5-4, PVI.V6-12, and PVI.V3-21, we wanted to map their epitopes on spike to better understand the molecular mechanisms of protection. We determined the cryo-EM structures of these mAbs (as Fabs to avoid cross-linking) in complex with a prefusion SARS-CoV-2/human/USA/USA-WA1/2020 HexaPro spike (Fig. 4). We obtained modest-resolution reconstruction for the PVI.V6-12 and PVI.V3-21 (Fig.4A, B) and a relatively high-resolution reconstruction for the PVI.V5-4 (Fig.4C-E) complexes. The structures confirm that PVI.V6-12 and PVI.V3-21 bind RBD; PVI.V6-12 most likely engages a complex epitope between two adjacent RBDs including the 360-370 α helix of one and the 500s loop of another RBD (Fig.4A). PVI.V6-12 likely does not discriminate between the “up” or “down” RBD conformations, whereas PVI.V3-21 requires an open RBD (Fig.4B). In fact, it appears that RBD is slightly dislodged “outward,” away from the 3-fold axis of the spike trimer, in our cryo-EM reconstruction, with PVI.V3-21 binding an epitope between the 340-helix and the 350-strand. While we note that due to the Fab occupancy and the particle alignment issues, we could not resolve the interaction interface on an individual amino acid level, for these two Fabs, we can conclude that both PVI.V6-12 and PVI.V3-21 still likely allow for the spike ACE2 receptor engagement and therefore do not neutralize the virus. Conversely, our 3.67Å resolution of the PVI.V5-4 Fab complex (Fig. 4C-E; S40) allowed us to perform a more detailed structural analysis of this complex. First, we discovered that the unknown epitope of PVI.V5-4 is the sub-domain 1 (SD1) within S1 (Fig. 4C), which is similar, but distinct from a recently reported sd1.040^32^. Second, our structure shows that one Fab is bound to one spike trimer and that the binding of PVI.V5-4 induces several structural rearrangements of the spike glycoprotein: (1) the RBD of the Fab-bound protomer is dislodged slightly more outward, away from the 3-fold axis of the trimer than in its canonically up conformation and (2) the Fab binding to one spike protomer likely induced the dissociation of the adjacent S1 domain, as only two S1 domains are present in our structure – indeed, in this conformation, PVI.V5-4 would sterically clash with the adjacent S1 NTD (Fig. S4). On the antibody side, PVI.V5-4 engages SD1 with both the heavy and the light chains (Fig.4D). Loops 2 and 3 of the PVI.V5-4 heavy chain complementarity determining region (HCDR) are sandwiching the SD1 550-560 loop. Additional intermolecular contacts are mediated by the LCDR3 loop (Fig.4D). The antibody HCDR3 Y111 and Y116, together with HCDR2 Y66 and LCDR3 Y92 amino acid side-chains, create an aromatic cradle that accommodates the spike SD1 150-160 loop (Fig.4E). We speculate that PVI.V5-4 binding still allows for ACE2 receptor engagement and hence does not neutralize the virus, but its epitope and angle of spike engagement position it to effectively engage FcR and provide protection. Hence, PVI.V5-4 is functionally distinct from sd1.040, which is a neutralizing mAb, by means of preventing ACE2-induced spike conformational changes^32^.

**Figure 4.**
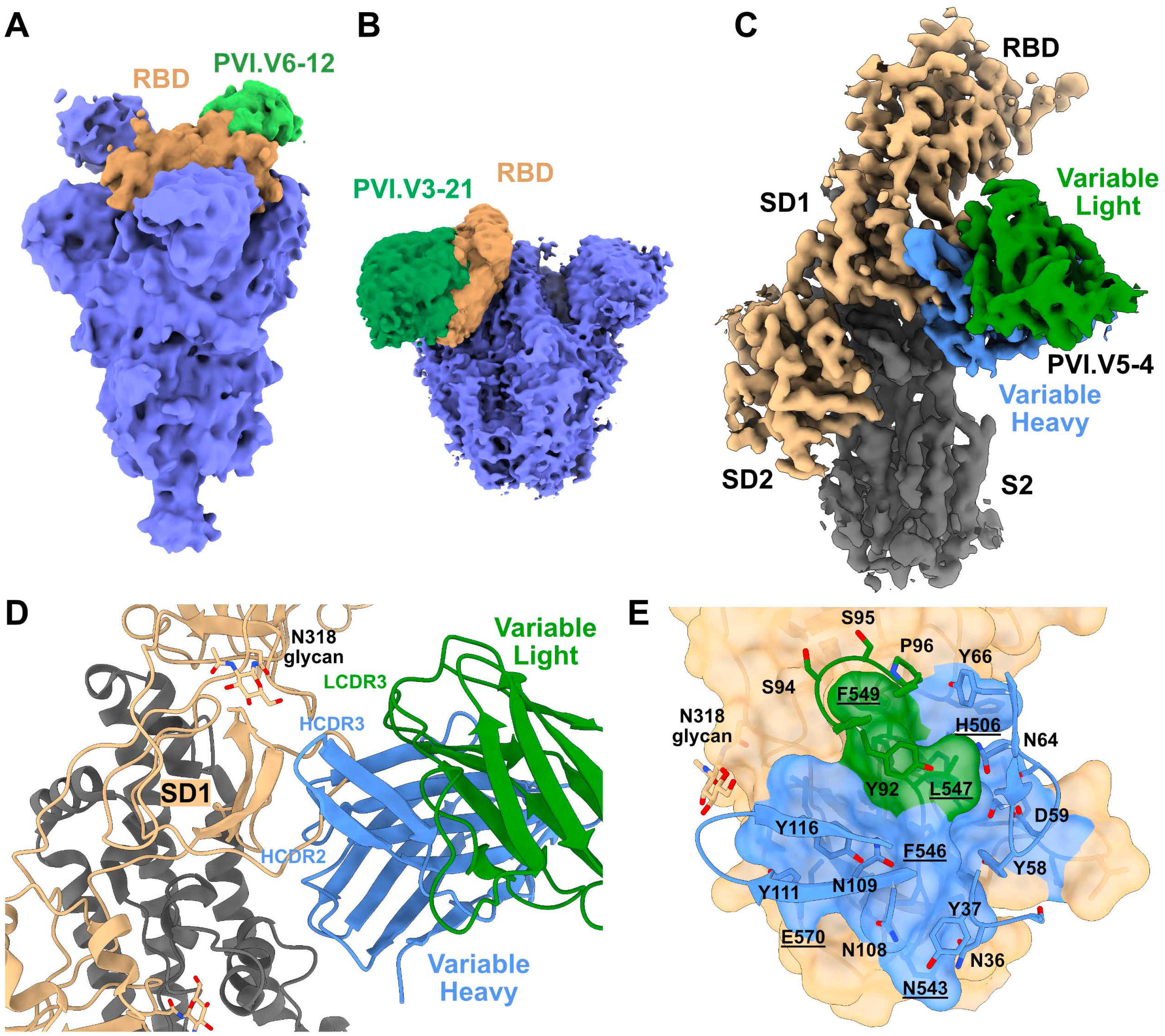
Cryo-EM structures of PVI.V6-12, PVI.V3-21, and PVI.V5-4 in complex with SARS-CoV-2 spike. (**A**) and (**B**) Global, modest-resolution cryo-EM maps of PVI.V6-12 and PVI.V3-21 Fabs in complex with spike. The structures identify the binding epitopes on the spike RBD. Fabs are colored green, RBD sienna, and the rest of spike blue. (**C**) Cryo-EM map of PVI.V5-4 in complex with spike at 3.67Å nominal resolution. The structure defines the PVI.V5-4 epitope on the sub-domain 1 (SD1). Spike S1 including RBD, SD1, and SD2 domains are colored sienna, S2 dark grey, and PVI.5-4 variable heavy and light chains blue and green, respectively. (**D**) Cartoon representation of the atomic model of PVI.V5-4 in complex with spike SD1. Major interactions are defined by the antibody HCDR3, HCDR2, and LCDR3 loops. N318 glycan is shown for context. € Surface representation of PVI.V5-4 epitope on spike SD1. The Fab interacting loops are shown as cartoon, and individual interacting residues are labeled and shown as sticks. The spike interacting residues are underlined. N318 glycan is shown for context.

### Non-neutralizing mAbs can mediate ADCC, ADCP, and complement binding via interactions with their Fc regions

To determine whether the mAbs elicit protection via effector functions, ADCC and ADCP reporter assays were performed using commercially available effector cells expressing either the human FcγRIIIa or FcγRIIa receptors respectively. The majority of the 42 mAbs tested exhibited FcγRIIIa activity in a luciferase reporter assay utilizing cells infected with recombinant Newcastle disease virus expressing the pre-fusion wild-type spike (NDV-HexaPro-S)^33^ (Fig. 5A). Neutralizing mAbs PVI.V3-9, PVI.V6-4, PVI.V6-2, PVI.V6-7, and PVI.V6-11 exhibited the highest activity. Of the non-neutralizing RBD binding mAbs, PVI.V6-3 exhibited the highest FcγRIIIa activity, while the protective mAbs PVI.V3-16, PVI.V3-21, PVI.V6-9, PVI.V6-12, and PVI.V6-15 exhibited comparable activity (Fig. 5A). Notably, mAbs PVI.V3-18, PVI.V3-19, and PVI.V6-5, which did not confer protection in the mouse challenge model, also demonstrated FcγRIIIa activity (Fig. 5A). For the NTD-binding mAbs, the protective mAb PVI.V6-8 exhibited FcγRIIIa activity, while the activity of PVI.V5-2 was lower. The non-protective mAb, PVI.V3-15, displayed comparable activity to PVI.V6-8, while mAbs PVI.V3-1 and PVI.V3-10 had little activity (Fig. 5A). The majority of the S2 binding mAbs displayed comparable FcγRIIIa activity, despite only about 30% of these providing protection from lethal challenge. Interestingly, the non-neutralizing mAbs PVI.V3-4, PVI.V5-2, and PVI.V5-4 did not display any FcγRIIIa activity despite their ability to confer survival and reduce lung viral titers in the mouse challenge experiments (Fig. 5A). Of the protective non-neutralizing mAbs, none demonstrated FcγRIIa activity, however, all the neutralizing mAbs, except for PVI.V3-9 and PVI.V6-4, successfully induced reporter assay responses (Fig. 5B). Overall, the majority of mAbs which were found to exhibit FcγRIIa binding activity were those targeting NTD.

**Figure 5.**
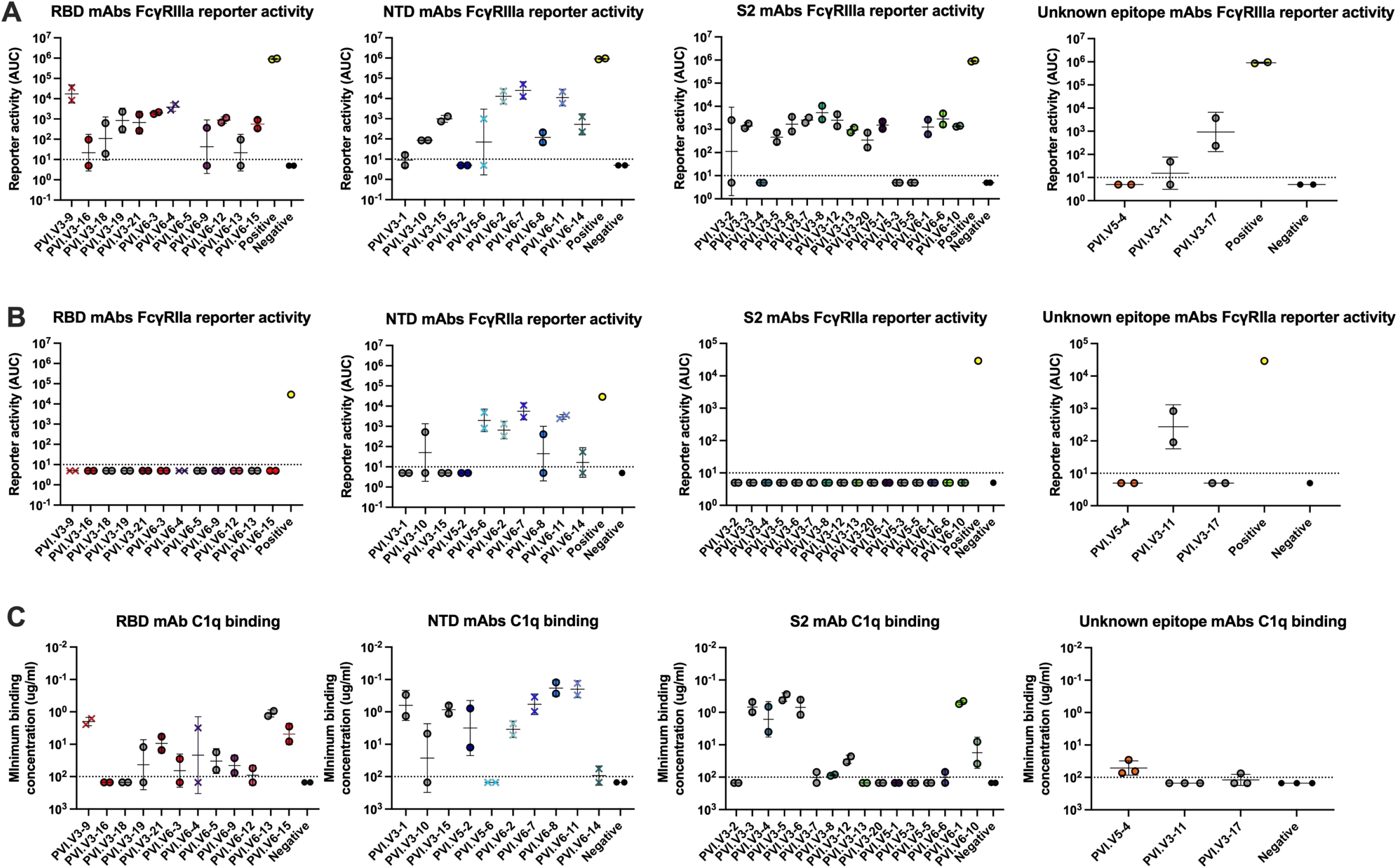
Characterization of the Fc-mediated activity of the mAbs. **(A)** FcγRIIIa reporter activity of the RBD, NTD, S2, and unknown epitope binding mAbs. **(B)** FcγRIIa reporter activity of the RBD, NTD, S2, and unknown epitope binding mAbs. **(C)** Binding of complement protein C1q to the Fc portion of the mAbs in the presence of wild type SARS-CoV-2 spike protein. Dotted lines denote the limit of detection.

To determine if the mAbs can bind the components of the complement pathway, enzyme-linked immunosorbent assays (ELISAs) were carried out in the presence of wild-type spike antigen, using secondary antibodies specific to human C1q. Of the RBD binding mAbs, the neutralizing mAbs PVI.V3-9 and PVI.V6-4 exhibited the highest C1q binding, as did the non-neutralizing protective mAbs PVI.V3-21, PVI.V6-3, PVI.V6-9, and PVI.V6-15 (Fig. 5C). Of the NTD binding, neutralizing mAbs, PVI.V6-11, PVI.V6-7 and PVI.V6-2 showed appreciable C1q binding, whereas PVI.V5-6 and PVI.V6-14 did not bind. Of the non-neutralizing NTD binding mAbs, all except PVI.V3-10 bound C1q (Fig. 5C). For the S2 binding mAbs, the protective, non-neutralizing mAbs PVI.V3-4, PVI.V6-1, and PVI.V6-10 displayed C1q binding, but PVI.V3-8, PVI.V3-13, PVI.V5-1, and PVI.V6-6 showed little to no binding (Fig. 5C). The non-protective, non-neutralizing mAbs PVI.V3-3, PVI.V3-5, and PVI.V3-6 also displayed appreciable C1q binding. Finally, the protective SD-1 binding mAb PVI.V5-4 exhibited weak, but detectable C1q binding (Fig. 5C).

### Some non-neutralizing antibodies retain their protective capabilities in the absence of effector functions

To confirm that the mAbs elicit their protective effects via Fc interactions, a subset of 6 non-neutralizing mAbs (PVI.V5-1, PVI.V5-2, PVI.V5-4, PVI.V6-3, PVI.V6-12, and PVI.V3-21) and one neutralizing mAb (PVI.V6-2) which conferred 100% survival against lethal challenge were cloned into IgG1 expression vectors that encode the LALA (L234A-L235A) or LALAPG (L234A-L235A, P329G) mutations^34,35^. The LALA mutation diminishes Fc binding by the FcγRs on effector cells and by complement, while the additional PG mutation abolishes all Fc binding^34,35^. ELISAs using ancestral SARS-CoV-2 spike confirmed that the binding of the LALA and LALAPG mAbs was comparable to their wild type counterparts (Fig. S3). These “effector silent” antibodies were utilized in ADCC/ADCP effector reporter assays in addition to C1q binding ELISAs. As expected, the LALA and LALAPG mutant mAbs no longer displayed FcγRIIIa, FcγRIIa reporter activity or C1q binding *in vitro* (Fig.6 A-C). The seven wild type mAbs were then employed in conjunction with their LALA and LALAPG counterparts in a challenge study in which mice were treated intraperitoneally with 10mg/kg of mAb and then infected with a 5xLD_50_ dose of SARS-CoV-2 MA10. Unsurprisingly, the sole neutralizing antibody, PVI.V6-2, maintained protection in the absence of effector functions at this concentration (Fig. 6D). Both the LALA and LALAPG mutant PVI.V5-2 were unable to protect mice from lethal challenge, with the mice succumbing to infection by 5dpi (Fig. 6D). The LALA and LALAPG mutant PVI.V3-21 antibodies displayed increased weight loss and decreased survival. Similarly, the LALA and LALAPG mutant PVI.V6-12 mAbs exhibited more weight loss than their wild-type counterpart, however the LALA mutant still conferred 100% survival. The non-neutralizing mAb PVI.V6-3 exhibited increased weight loss in the presence of the LALAPG mutations, however the LALA mutant mAb promoted protection compared to wild-type. Surprisingly, antibodies PVI.V5-4 and PVI.V5-1 retained protection even in the absence of effector functions. As these LALA and LALAPG mutant mAbs do not neutralize SARS-CoV-2, and lack both effector cell and complement binding, the mechanism through which they promote survival *in vivo* is unclear.

**Figure 6.**
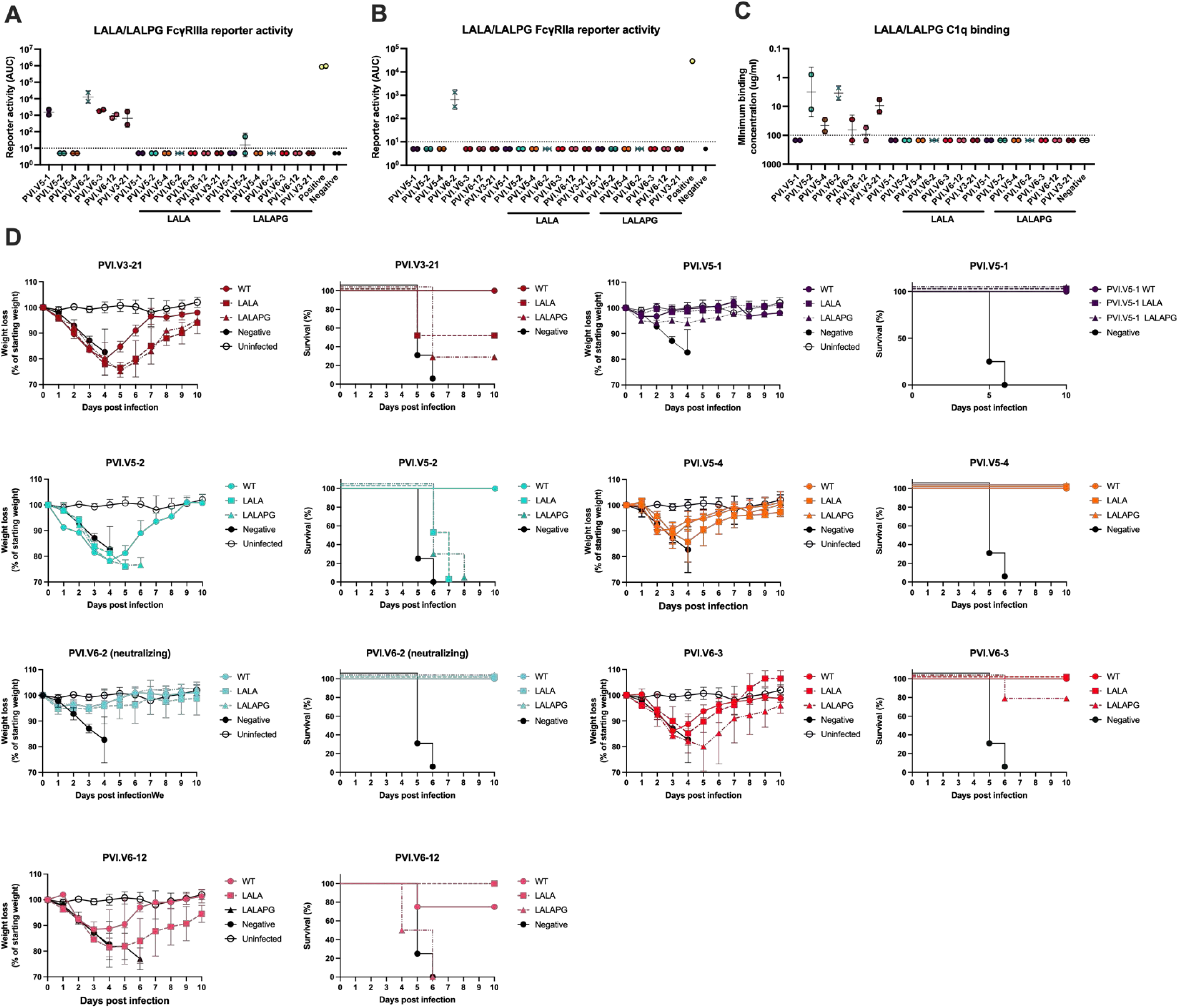
Investigating the protection elicited by Fc-silent mutant mAbs. **(A)** FcγRIIIa reporter activity of the seven most protective mAbs and their LALA and LALAPG mutant counterparts. **(B)** FcγRIIa reporter activity of the wild type, LALA, and LALAPG mAbs. **(C)** Binding of complement protein C1q by the wild type, LALA, and LALAPG mAbs in the presence of wild type SARS-CoV-2 spike protein. Dotted lines denote the limit of detection. **(D)** Weight loss and survival of BALB/cAnNHsd mice treated the seven most protective mAbs and their LALA and LALAPG counterparts following lethal challenge with mouse-adapted SARS-CoV-2.

## Discussion

Four years after its emergence, SARS-CoV-2 remains a serious threat to human health. With the continued emergence of immune evasive SARS-CoV-2 VOCs that harbor spike mutations capable of ablating neutralizing antibodies, the mechanisms that maintain protection without these responses require further characterization. Numerous studies have shown that neutralizing antibodies elicited by vaccination and/or infection correlate with protection from SARS-CoV-2 infection^12–14,36–39^. Conversely, a wealth of literature that indicates that antibody functions other than direct neutralization also contribute to protection from disease^40–49^. Additionally, several studies have shown that, while SARS-CoV-2 variants are able to evade the neutralizing antibody response, conserved epitopes are still recognized by CD4+ and CD8^+^ T cells induced by both vaccination and infection^50–54^. In a previous study, we demonstrated that SARS-CoV-2 elicits a higher proportion of non-neutralizing antibodies versus neutralizing antibodies, predominantly against regions of the spike protein outside of the RBD^10^. To determine whether these mAbs contribute to protection against SARS-CoV-2, we studied the ability of these mAbs to protect mice from lethal challenge with mouse-adapted SARS-CoV-2. Seven of the 42 mAbs have previously been demonstrated to neutralize the virus, while the remainder do not. As expected, all the neutralizing mAbs protected mice against lethal challenge, however, with variable potency. Of the non-neutralizing mAbs, 19 promoted survival following lethal challenge, most of which were previously found to bind within the RBD. While most of the tested mAbs bind within S2, only 5 of these were found to be protective. We further validated the protective capabilities of these mAbs by testing the six most protective non-neutralizing mAbs and one neutralizing mAb using the hamster and hACE2 K-18 challenge models. These experiments confirmed that non-neutralizing mAbs can promote protection from challenge with ancestral SARS-CoV-2, with some mAbs exhibiting comparable protection to neutralizing mAbs. Furthermore, these mAbs showed protection even when tested using highly lethal challenge doses in animal models of SARS-CoV-2 infection which display severe disease phenotypes. While these mAbs cannot block the virus, their binding to cells which display spike on their surface resulted in the engagement of Fc-medicated ADCC in a reporter assay.

During natural infection, the Fc portion of antibodies bound to virions and/or infected cells can interact with the FcγRs of a diverse array of immune cells. The FcγRIIIa receptor is the sole FcγR expressed on NK cells^50^ that, when engaged, initiates a signaling cascade involving the phosphorylation of immunoreceptor tyrosine-based inhibitory motifs (ITIMs)^55–58^ thereby promoting ADCC activity. Similarly, the FcγRs expressed on macrophages, monocytes, and granulocytes promote ADCP when interacting with the Fc portions of antibodies bound to the viral glycoproteins present on virions and infected cells. Through these mechanisms, non-neutralizing antibodies can promote the clearance of virions and infected cells and, as a result, curtail infection and reduce viral loads in infected tissues. The Fc regions of antibodies bound to free virions can also be engaged by C1q, the first protein involved in the classical complement cascade. The complement system is composed of over 30 heat-liable proteins that are present within plasma^21,59–61^. The engagement of the Fc region of antibodies by Cq1 initiates an enzymatic cascade which generates the membrane attack complex (MAC), a pore-forming complex that ruptures lipid bilayers. Importantly, the MAC can form on infected cells and virus particles, providing a mechanism through which virus-producing cells and free virions can be destroyed.

Most of the non-neutralizing mAbs described herein can engage both effector cells and C1q via their Fc domains, indicating that this is likely the mechanism through which they protect mice from lethal challenge. Indeed, at 3 dpi, the neutralizing mAbs targeting the RBD had effectively reduced viral loads below the limit of detection, whereas the non-neutralizing antibodies reduced loads to a lesser, but still significant extent. Of the five NTD-binding neutralizing mAbs, only PVI.V6-11 reduced viral loads below the limit of detection at 3dpi, however PVI.V6-2 and PVI.V6-7 both promoted protection from lethal infection despite maintaining high lung titers. The mice treated with the majority of the non-neutralizing mAbs exhibited increased weight loss and lung viral loads compared to those treated with neutralizing mAbs, demonstrating that non-neutralizing mAbs are less efficient at restricting viral infection. Given that neutralizing mAbs are capable of inactivating free virions, and non-neutralizing mAbs predominantly work upon infected cells, these results indicate that non-neutralizing antibodies can still elicit effective protection from disease, however not to the same extent as neutralizing antibodies – and from a mechanistic perspective they cannot protect from infection. Conversely, as the majority of non-neutralizing mAbs tested here retain binding to variant SARS-CoV-2 spikes, the effector functions may help ameliorate disease burden upon infection with SARS-CoV-2 VOCs.

To test whether the most protective non-neutralizing antibodies restrict viral infection via Fc medicated interactions, six of the non-neutralizing mAbs which conferred 100% survival against lethal challenge were cloned into expression vectors harboring the LALA or LALAPG mutations in their Fc regions^62,63^. One protective, RBD-binding, neutralizing mAb, PVI.V6-2, was also mutagenized as a control. These mutations have been used extensively in previous research wherein mAbs have been developed for therapeutic use^34,35,63–66^. ADCC and ADCP effector cell reporter assays and complement ELISAs confirmed that the inclusion of both the LALA and LALAPG mutations abrogated the effector functions of these mAbs. When this panel of mAbs was investigated in the mouse challenge model, as expected, the neutralizing mAb PVI.V6-2 retained 100% protection, even in the presence of the LALA and LALAPG mutations. While neutralizing mAbs can also engage effector cells and complement, the neutralization of virus is the most likely mechanism through which this mAb protects against infection. Conversely, both the LALA and LALAPG counterparts of the non-neutralizing mAb PVI.V5-2 no longer protected mice from infection, indicating that it likely interacts predominantly with effector cells to promote ADCC and ADCP. Similarly, the non-neutralizing mAb PVI.V3-21 LALA mutant elicited 50% protection, whereas the LALAPG mutant provided 20% protection. This indicates that this mAb likely engages both complement and effector cells. The non-neutralizing mAb PVI.V6-12 lost all protective capabilities following LALAPG mutagenesis, however provided 100% protection from infection with only LALA. Surprisingly, the non-neutralizing antibodies PVI.V5-1, PVI.V6-3, and PVI.V5-4 retained protection even when mutagenized to include the LALA and LALAPG substitutions. PVI.V6-3 exhibited weak *in vitro* neutralizing activity and is likely incapable of neutralizing virus at the 10 mg/kg dose used, however this activity may explain some of its protective effects in the presence of the LALA and LALAPG mutations. As PVI.V5-1 and PVI.V5-4 are unable to neutralize and, when mutagenized, longer engage effector cells or complement, the mechanism through which they induce protection from infection is unclear. It has previously been reported that non-neutralizing Ebola virus mAbs, mutagenized with the G236R/L328R mutation which abrogates Fc binding, retained protection in a mouse challenge model^67^. This indicates that the mechanisms through which these mAbs inhibit virus cannot be recapitulated using *in vitro* neutralization assays and can only be probed using animal models.

Structural epitope mapping can provide invaluable information about the molecular mechanisms of antibody protection. We used single-particle cryo-EM to glean insights into how non-neutralizing antibodies protect by virtue of engaging the spike on infected cells, activating phagocytic and cytolytic cells and thus helping reduce the disease burden in infected individuals. We showed that PVI.V3-21, PVI.V5-4, and PVI.V6-12 bind epitopes that do not interfere with the spike cellular receptor ACE2 binding but potentially position their Fc portion for efficient activation of effector functions like ADCC, ADCP, and complement activation. Notably, we also report on a detailed structural characterization of a non-neutralizing human antibody that targets SD1, a vastly underappreciated yet conserved domain on spike adjacent to RBD. Previous studies have shown that interactions between the RBD and SD1 of one spike S1 protomer and the NTD of the neighboring S1 protomer can stabilize the spike in its prefusion conformation^68^. Our structure of PVI.V5-4 bound to SD1 shows that antibodies that engage SD1 destabilize those protomer-protomer interactions and dislodge the neighboring NTD. Thus, while PVI.5-4 is devoid of neutralizing and effector functions, it does prevent weight loss in a murine model of infection, probably by destabilizing the spike quaternary architecture on infected cells. While PVI.5-4 showed no neutralizing activity in our *in vitro* assays, other SD1-targeting antibodies have been shown to neutralize the virus by inhibiting the conformational changes that would expose the S2 fusion loops upon ACE2 receptor engagement^32^. We believe that SD1-targeting antibodies could be a valuable component of the humoral response against SARS-CoV-2 and other coronaviruses. Questions remain about their prevalence in infection- and vaccination-induced humoral immunity.

Taken together, these results suggest that non-neutralizing antibodies play important roles in the protecting against COVID-19, that such mAbs may provide protection against emerging variants and, if sufficiently cross-reactive, perhaps against other betacoronaviruses^69^. Critically, while some non-neutralizing SARS-CoV-2 mAbs have been reported to exhibit antibody-dependent enhancement (ADE) *in vitro*^70–73^, our study and others demonstrate a lack of ADE *in vivo*^72^ indicating that mAbs with no neutralizing activity do not promote enhanced disease phenotypes. Finally, our findings are in line with reports that show binding antibody levels correlation with protection from severe outcomes in humans in the absence of robust neutralizing activity^17^.

### Limitations of the study

Some important caveats of this work should be considered. As we have performed effector cell reporter assays using recombinant cells which express either hFcγRIIIA or hFcγRIIA receptors, we are unable to model the complex expression pattern of these receptors which are present together on some immune cell populations and function in conjunction with the inhibitory receptor FcγRIIB. Additionally, there are important differences between the human and murine FcγRs, wherein mice express four different, but homologous receptors (mFcγRI, mFcγRIIB, mFcγRIII and mFcγRIV). Despite these complexities, human IgG can bind orthologous murine and hamster FcγRs with similar affinity to their human counterparts^74,75^, however mice do not recapitulate the diversity or expression patterns of the human FcγRs^76^. In future studies, these differences could be overcome by using humanized mice which express the human FcγRs^77^. Additionally, we have performed challenge studies using a single 10 mg/kg dose of mAb; differences in weight loss and survival may be more obvious when utilizing de-escalating doses.

## Material and methods

### Cells and viruses

Vero.E6 cells (American Type Culture Collection, ATCC, CRL-1586, clone E6) were maintained in Dulbecco’s modified Eagle medium (DMEM; Gibco), supplemented with Antibiotic-Antimycotic (100 U/mL penicillin– 100 μg/mL streptomycin–0.25 μg/mL amphotericin B; Gibco), and 10% fetal bovine serum (FBS; Corning) in a 37°C incubator with 5% CO_2_. Vero.E6 cells expressing Transmembrane protease, serine 2 (TMPRSS2, BPS Biosciences, 78081) were cultured in Dulbecco’s modified Eagle medium (DMEM; Gibco) supplemented with 10% FBS, 1% minimum essential medium (MEM) amino acids solution (Gibco, cat. no. 11130051), 100 U ml−1 penicillin and 100 μg mL−1 streptomycin (Gibco, cat. no. 15140122), 100 μg mL−1 normocin (InvivoGen, cat. no. ant-nr), and 3 μg mL−1 puromycin (InvivoGen, cat. no. ant-pr). Expi293F cells were maintained in Expi293 media in a shaking incubator at 37°C and 8% CO_2_.

Mouse-adapted SARS-CoV-2 (SARS-CoV-2 MA10, BEI Resources NR-55329), wild-type SARS-CoV-2 (isolate USA-WA1/2020), and Beta SARS-CoV-2 (B.1.351, hCoV-19/South Africa/ KRISP-K005325/2020, BEI Resources NR-54009), were obtained from BEI Resources. SARS-CoV-2 BA.1 (B.1.1.529, USA/NY-MSHSPSP-PV44488/2021, EPI_ISL_7908059) and SARS-CoV-2 BA.5 USA/NY-MSHSPSP-PV58128/2022 have been described previously^78^. Wild-type SARS-CoV-2 and SARS-CoV-2 MA10 were grown in Vero.E6 cells whereas Beta, Delta, BA.1, and BA.5 SARS-CoV-2 were grown using Vero.E6 TMPRSS2 cells. All viruses were grown for 3 days at 37°C prior to clarification via centrifugation at 1,000g for 10 minutes. Viral stocks used were sequence-verified and tittered using both the 50% tissue culture infectious dose (TCID_50_) method and via plaque assay on Vero.E6 TMPRSS2 cells before use in microneutralization assays and challenge experiments. Virus stocks were stored at −80°C before use. All experiments with authentic replication competent SARS-CoV-2 were performed in the biosafety level 3 (BSL-3) facility following institutional guidelines.

### Recombinant protein production

Recombinant proteins were expressed in Expi293F cells (Life technologies). Full-length spike proteins were cloned into the pCAGGS mammalian expression vector as previously described^79,80^. MAbs were also produced via the transfection of expression vectors encoding the mAb heavy chain and light chains as previously described^10^. Cells were grown to 3×10^6^ cells/mL and transfected using Expifectamine (Life Technologies) with 1 μg of plasmid per mL of culture media, as per manufacturer’s instructions. Enhancer solutions were added 24 hours post-transfection and cells were allowed to incubate for 3 days. At this point, the supernatant was removed, clarified by centrifugation at 4,000 g for 30 mins, and filtered using 0.45 μM Stericup filters (Sigma-Aldrich). Proteins were purified from filtered supernatants using nickel nitriloacetic acid (Ni-NTA) agarose (QIAGEN) via gravity flow, and proteins were eluted in line with previous publications^10^. Eluted proteins were buffer exchanged using 50 kDa Amicon centrifugal units (EMD Millipore), resuspended in phosphate-buffered saline (PBS), and quantified using a Bradford assay (Bio-Rad). Human IgG1 expression vectors harboring the LALA and LALAPG mutations were kindly provided by Ali Ellebedy. The VH genes of PVI.V3-21, PVI.V5-1, PVI.V5-2, PVI.V5-4, PVI.V6-2, PVI.V6-3, and PVI.V-12 were amplified via PCR and cloned into the LALA and LALAPG IgG1 vectors using InFusion cloning (Takara).

### ELISA

Recombinant proteins were coated on Immulon 4 HBX 96-well plates (Thermo Scientific) at a concentration of 2 μg/mL in PBS, 50 μL per well overnight at 4°C. 24 hours later, the coating solution was removed, and plates were blocked with 3% non-fat milk (Life Technologies) diluted in PBS containing 0.01% Tween-20 (PBST; Fisher Scientific). Plates were incubated with the blocking solution for 1 hour prior to 3 washes using PBST. MAbs were serially diluted 1:10 from a starting concentration of 30 μg/mL in PBST supplemented with 1% non-fat milk and were added to the respective plates. Primary antibody was not added to 16 wells per ELISA plate to serve as blank. After a 1-hour incubation at room temperature, the plates were washed 3 times with PBST and 100 μL of mouse anti-human IgG conjugated to horseradish peroxidase (HRP) diluted 1:3000 in PBST supplemented with 1% non-fat milk was added to each well. After an incubation time of 1 hour at room temperature, plates were again washed three times with PBST prior to the addition of 100 μL of o-phenylenediamine dihydrochloride (Sigma-Aldrich; OPD). Plates were allowed to develop for 10 minutes after which the reaction was stopped via the addition of 50 μL of 3M hydrochloric acid (HCl). The plates were read using a Synergy 4 (BioTek) plate reader at an optical density of 490 nanometers. The area under the curve (AUC) and the minimal binding concentrations for each plate was calculated using GraphPad Prism 10, with the average plus 5 standard deviations of the blank wells serving as the baseline. C1q and C3 binding ELISAs were carried out as previously described. Briefly, mAbs were diluted to 100 µg/mL and serially diluted 1:3 in sterile PBS. 50 µL of these dilutions were utilized to coat Immulon 4 HBX 96-well plates (Thermo Scientific) overnight at 4°C. The plates were then washed three times with PBST and blocked using 100 µL of protein-free blocking buffer (Pierce) per well for 1 hour. Human serum was then diluted to 3% (vol/vol) in protein-free blocking buffer (Pierce), and 100 µL was incubated per well for 1 hour. Following an additional three PBST washes, 100 µL of biotinylated anti-human C1q (ab48341, Abcam) or anti-human C3 (ab48342, Abcam), diluted 1:3000 in protein-free blocking buffer was added to each well and incubated for 1 hour at room temperature. Following three PBST washes, 100 µL of high sensitivity streptavidin conjugated to HRP (Thermofisher) diluted 1:3000 in protein-free blocking buffer was added. The plates were developed using an identical method to the conventional ELISA, and the data was analyzed using GraphPad Prism 10.

### Microneutralization assays

All work with SARS-CoV-2 was performed in a biosafety level 3 (BSL3) laboratory. Vero.E6 TMPRSS2 cells were seeded in a 96-well cell culture plate (Corning; 3340) at a density of 1×10^4^ cells per well. MAb dilutions were prepared in 1× minimal essential medium (MEM; Gibco) supplemented with 2% FBS. The irrelevant influenza A (IAV) antibody, CR9114, was included as a negative control^48^. 24 hours later cells were infected with 10^4^ tissue culture infectious dose 50 (TCID_50_)/mL, and 80 μL of virus and 80 μL of antibody dilution were combined and incubated together for 1 hour at room temperature on a separate 96-well plate. After the incubation, 120 μL of virus–antibody inoculum was utilized to infect cells for 1 hour at 37°C. After one hour, the inoculum was removed and 100 μL of each corresponding dilution was added to the wells. A volume of 100 μL of 1X MEM was also added to the plates to achieve a total volume of 200 μL per well. Cells were incubated at 37°C in a 5% CO_2_ incubator for 3 days and fixed with 10% paraformaldehyde (Polysciences) for 24 hours. The paraformaldehyde was removed, and cells were washed twice with PBS prior to permeabilization by the addition of 150 µL of 0.1% Triton X-100 (Fisher followed by incubation at room temperature for 15 minutes. The Triton X-100 solution was removed, cells were blocked with PBS supplemented with 3% non-fat milk (American Bio; catalog no. AB10109-01000) for one hour, and then stained using the anti-nucleoprotein antibody 1C7 as previously described^81^.

### ADCC and ADCP *in vitro* reporter assays

Vero.E6 cells (ref) were seeded in 96 well plates (Corning; 3340) at a seeding density of 2×10^4^ cells per well 24 hours prior to infection with NDV-HexaPro-S^82^ at a multiplicity of infection (MOI) of 2. One hour later the inoculum was removed and replaced with 100µl of Roswell Park Memorial Institute (RPMI-1640) media (Thermofisher). 48 hours later the media was removed and replaced with 25 µL of RMPI-1640 media supplemented with 2% low-IgG FBS (Promega). Antibodies were serially diluted 1:10 in RPMI 1640 media from a starting concentration of 30 µg/mL to a final concentration of 0.014 µg/mL and 25 µL of antibody dilution was added to the plate. CR9114 was utilized as a negative control^48^. MAb dilutions were not added to 16 wells per plate to provide background luminescence readings. 3×10^6^ ADCC and ADCP Bioassay Effector Cells expressing the human FcγRIIIa or FcγRIIa receptors (Promega) were added per well and the cells were incubated in at 37°C with 5% CO_2_ for 6 hours. 75 µL of Bio-Glo luciferase reagent (Promega) was then added to each well and the luminescence was read in a Synergy 4 (BioTek) plate reader. The background luminescence reading was used to calculate AUC using GraphPad Prism 10.

### In vivo studies

All animal procedures were performed by following the Institutional Animal Care and Use Committee (IACUC) guidelines of the Icahn School of Medicine at Mount Sinai IACUC and according to an approved protocol. To determine the LD_50_ of SARS-CoV-2 MA10, 6- to 8-week-old female, BALB/cAnNHsd mice (Envigo) were infected with serial dilutions of the virus ranging from 10^5^ to 5 plaque forming units (PFU). LD50 experiments were carried out identically in the hACE2 K-18 mice using SARS-CoV-2/human/USA/USA-WA1/2020. For antibody protection studies, 6–8-week-old female BALB/cAnNHsd or B6.Cg-Tg(K18-ACE2)2Prlmn/J mice (The Jackson Laboratory) were treated intraperitoneally with 10 mg/kg of antibody diluted in sterile PBS. Two hours post-treatment mice were administered anesthesia intraperitoneally (IP) and intranasally (IN) infected with 50 μL of either 1xLD_50_ or 5xLD_50_ of SARS-CoV-2 MA10 or WA1/2020 diluted in sterile PBS. Anesthesia was prepared using 0.15 mg/kg ketamine and 0.03 mg/kg xylazine in water for injection (WFI; Gibco). Weights were recorded over a period of 1-2 weeks and mice falling below 75% of their original body weight were humanely sacrificed via cervical dislocation. When assessing lung viral loads, mice were humanely sacrificed on days 3- and 5-dpi for enumeration of virus in the lungs. Lungs were homogenized using a BeadBlaster 24 (Benchmark) homogenizer and clarified by centrifugation at 1,000 g for 10 minutes at 4°C. Viral load in the lung was quantified via a classic plaque assay. For the hamster studies, 6-8-week-old female Syrian hamsters (Envigo) were treated intraperitoneally with 10 mg/kg of antibody diluted in sterile PBS. Two hours later, hamsters were anesthetized via the IP administration of 50-200 mg/kg ketamine and 5-10 mg/kg xylazine and infected IN with 100 µL sterile PBS containing 10^5^ PFU of wild-type SARS-CoV-2 (isolate USA-WA1/2020). Weight loss was recorded for 10 days. The IAV mAb, CR9114, was included as a negative control in all animal studies^48^.

### Cryo-EM sample preparation and data collection

SARS-CoV-2 HexaPro Spike^83^ was incubated with PVI.V3-21, PVI.V5-4, or PVI.V6-12 Fabs at 2 mg/mL and a molar ratio of 1.2:1 Fab:Spike for 20 minutes at 4°C. Immediately before grid preparation, fluorinated octyl maltoside was added to the pre-formed complex at 0.02% w/v final concentration. Then, 3 µL aliquots were applied to UltrAuFoil gold R0.6/1 grids (Quantifoil) and subsequently blotted for 6 seconds at blot force 0, then plunge-frozen in liquid ethane using an FEI Vitrobot Mark IV. Grids were imaged on a Titan Krios microscope operated at 300 kV and equipped with a Gatan K3 Summit direct detector. 5,894 movies were collected in counting mode at a total dose of 52.51 e^-^/Å^2^ at a magnification of 81,000, corresponding to a calibrated pixel size of 1.069 Å. Defocus values were from −0.5 to −1.5 μm.

### Cryo-EM data processing

Movies were aligned and dose weighted using MotionCorr2^84^. Contrast transfer function estimation was done in cryoSPARC v3.3.1^85^ using CTFFIND4 and particles were picked with cryoSPARC’s blob picker. The picked particles were extracted with a box size of 512 pixels, with 4x binning and subjected to a 2D classification. Selected particles were then subjected to a second round of 2D classification. An initial model was generated on the 951,973 selected particles at 6 Å/pixel. After one round of non-uniform refinement, without imposed symmetry, the particles were subjected to 3D classification with 10 classes. Of these, the best 3 classes, containing 173,783 particles in total, were combined, re-extracted without binning with a box size of 512 pixels, and selected for further rounds of non-uniform refinement with local CTF refinement, yielding the final global map at a nominal resolution of 3.35 Å. The protomer with the Fab volume was subjected to local refinement with a soft mask extended by 6 pixels and padded by 12 pixels encompassing the RBD, SD1, and Fab. This yielded the final local map at 3.67 Å resolution. The two half-maps from the global or local refinement were used for sharpening in DeepEMhancer (PMID: 34267316) The reported resolutions are based on the gold-standard Fourier shell correlation of 0.143 criterion.

### Model building and refinement

Spike from PDB ID: 6ZGE and SAbPred^86^ generated Fab variable domains were fit into the focus refined map using UCSF ChimeraX^87^ and then manually built using COOT^88^. N-linked glycans were built manually in COOT using the glyco extension and their stereochemistry and fit to the map validated with Privateer^89^. The model was then refined in Phenix^90^ using real-space refinement and validated with MolProbity^91^. The structural biology software was compiled and distributed by SBGrid^92^.

## Data availability

The atomic models and corresponding maps are available online under the following accession codes. For the PVI.V5-4 complex, the atomic model is available in the Protein Data Bank (PDB) under accession code 8VIA. The cryo-EM maps and the raw micrographs are available in the Electron Microscopy Data Bank (EMDB) and in the Electron Microscopy Public Image Archive (EMPIAR) under accession codes EMD-43255 and EMPIAR-11847, respectively. Additionally, cryo-EM maps for the PVI.V3-21 and PVI.V6-12 complexes have been deposited to EMDB under accession codes EMD-43306 and EMD-43314, respectively. All other data underlying figures in this paper can be found in ImmPort under identifier xxx.

## Acknowledgments

We thank the Mount Sinai Pathogen Surveillance Program for providing representative SARS-CoV-2 viral isolates.

G.B. was in part supported by NIH R01AI168178 and the Irma T. Hirschl/Monique Weill-Caulier Trust.

Some of this work was performed at the National Center for CryoEM Access and Training (NCCAT) and the Simons Electron Microscopy Center located at the New York Structural Biology Center, supported by the NIH Common Fund Transformative High Resolution Cryo-Electron Microscopy program (U24 GM129539), and by grants from the Simons Foundation (SF349247) and NY State Assembly.

This effort was supported by the Serological Sciences Network (SeroNet) in part with Federal funds from the National Cancer Institute, National Institutes of Health, under Contract No. 75N91019D00024, Task Order No. 75N91021F00001. The content of this publication does not necessarily reflect the views or policies of the Department of Health and Human Services, nor does mention of trade names, commercial products or organizations imply endorsement by the U.S. Government.

This work was also partially funded by the Centers of Excellence for Influenza Research and Response (CEIRR, contract # 75N93021C00014, reagent generation), by the Collaborative Influenza Vaccine Innovation Centers (CIVICs contract # 75N93019C00051, reagent generation), P01 AI172531 (PLUTO), U19AI168631 (VIVA) and by institutional funds.

This work was supported in part through the computational and data resources and staff expertise provided by Scientific Computing and Data at the Icahn School of Medicine at Mount Sinai and supported by the Clinical and Translational Science Awards (CTSA) grant UL1TR004419 from the National Center for Advancing Translational Sciences. Research reported in this publication was also supported by the Office of Research Infrastructure of the National Institutes of Health under award number S10OD026880 and S10OD030463. The content is solely the responsibility of the authors and does not necessarily represent the official views of the National Institutes of Health.

This work was partially supported by the São Paulo Research Foundation (R.A-S fellowship), projects 2023/02345-7, 2021/05661-1 and 2020/08943-5.

---

**Figure S1.**
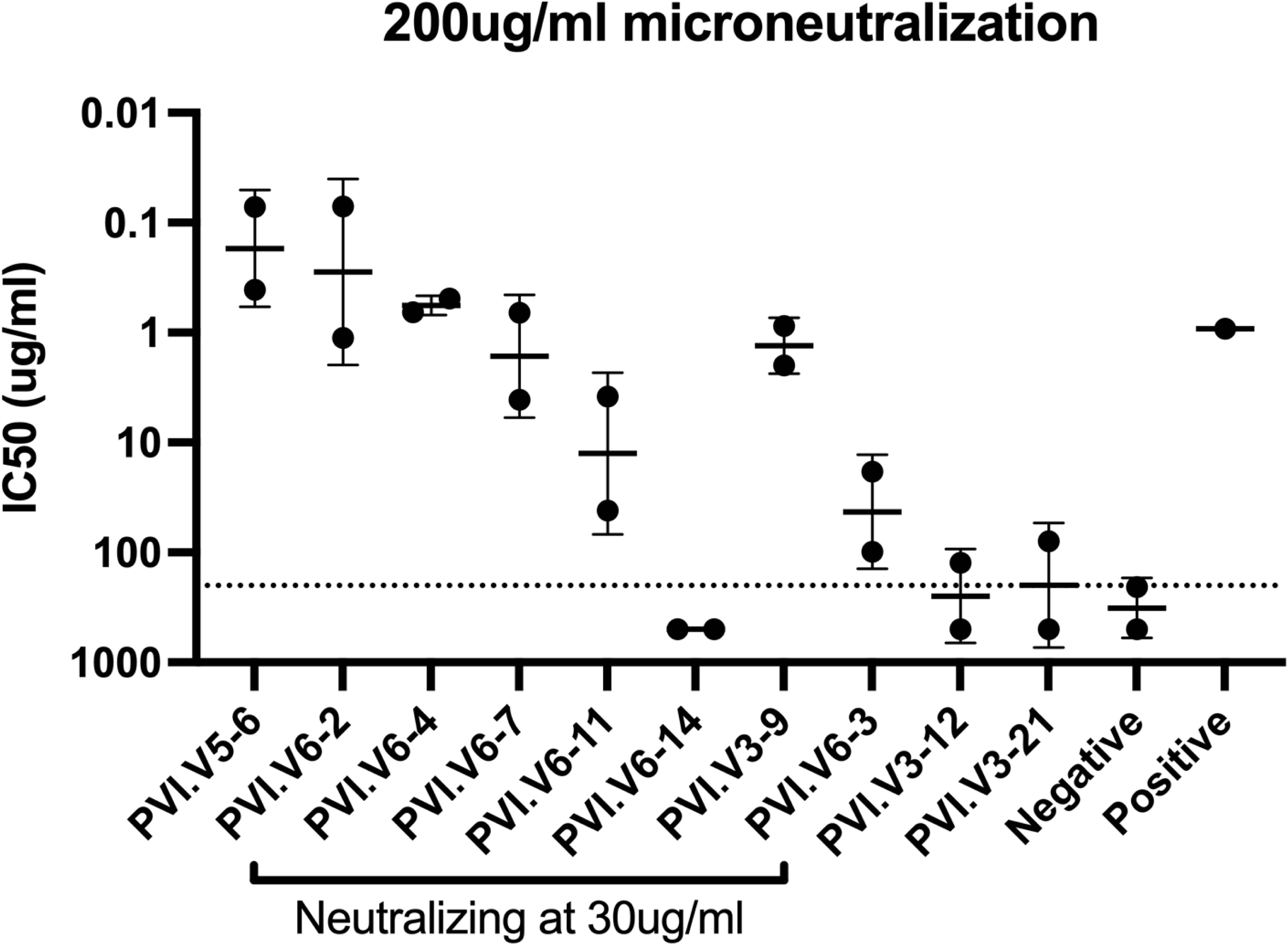
Neutralizing activity of the mAbs at high concentration. Neutralizing activity of the mAbs against authentic SARS-CoV-2 at a starting concentration of 200 μg/ml. Only mAbs exhibiting detectable neutralization are shown.

**Figure S2.**
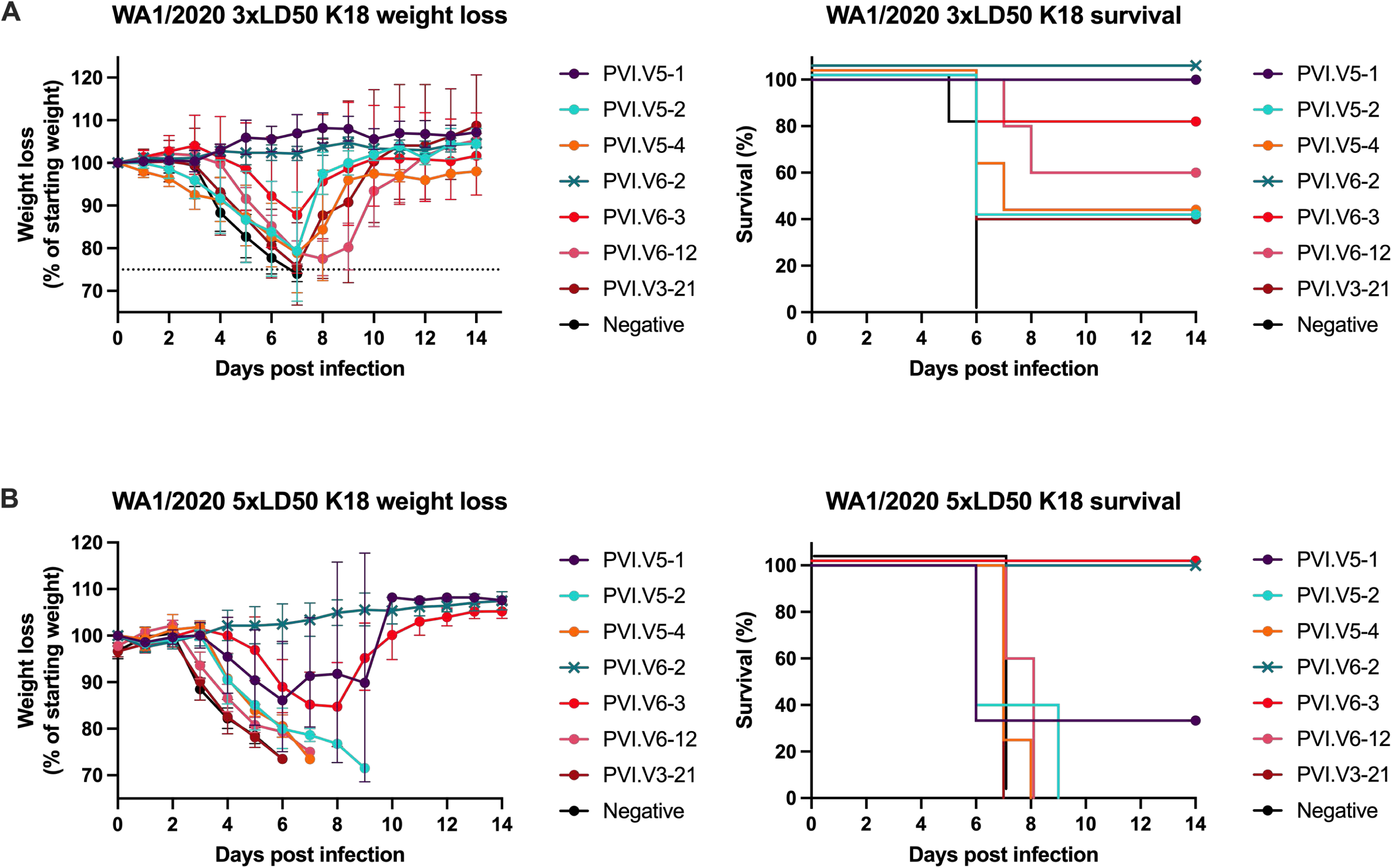
Protection elicited by the 7 most protective mAbs in the K18 murine model. **(A)** Weight loss and survival in transgenic K18 mice following treatment with mAbs and challenge with a 3x LD_50_ dose of wild type SARS-CoV-2. **(B)** Weight loss and survival in K18 mice following treatment with mAbs and challenge with a 5x LD_50_ dose of wild type SARS-CoV-2.

**Figure S3.**
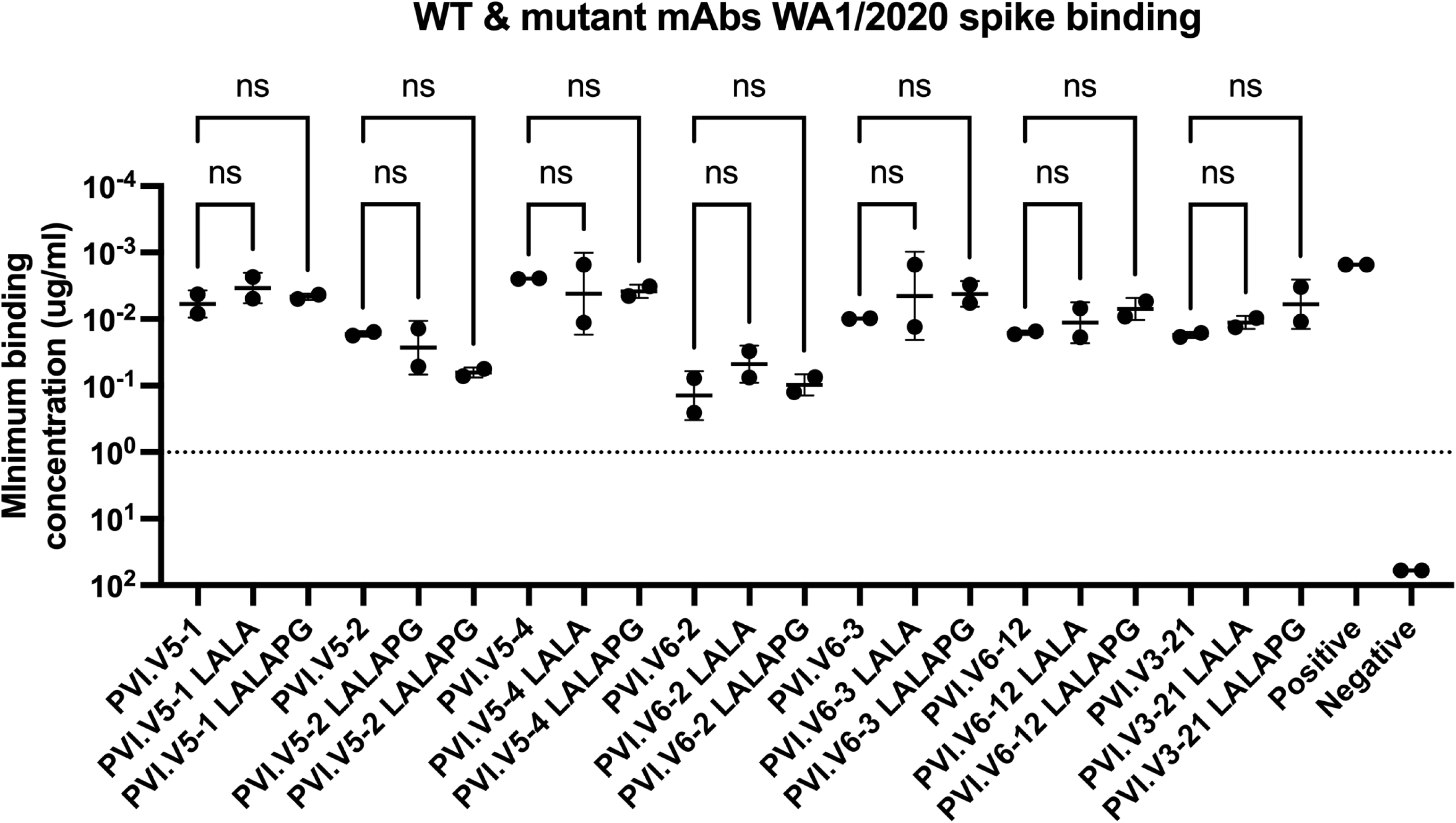
Binding of wild type, LALA, and LALAPG mAbs to SARS-CoV-2 spike. Binding of the seven most protective SARS-CoV-2 mAbs and their mutant LALA and LALAPG equivalents to wild type SARS-CoV-2 spike.

**Figure S4.**
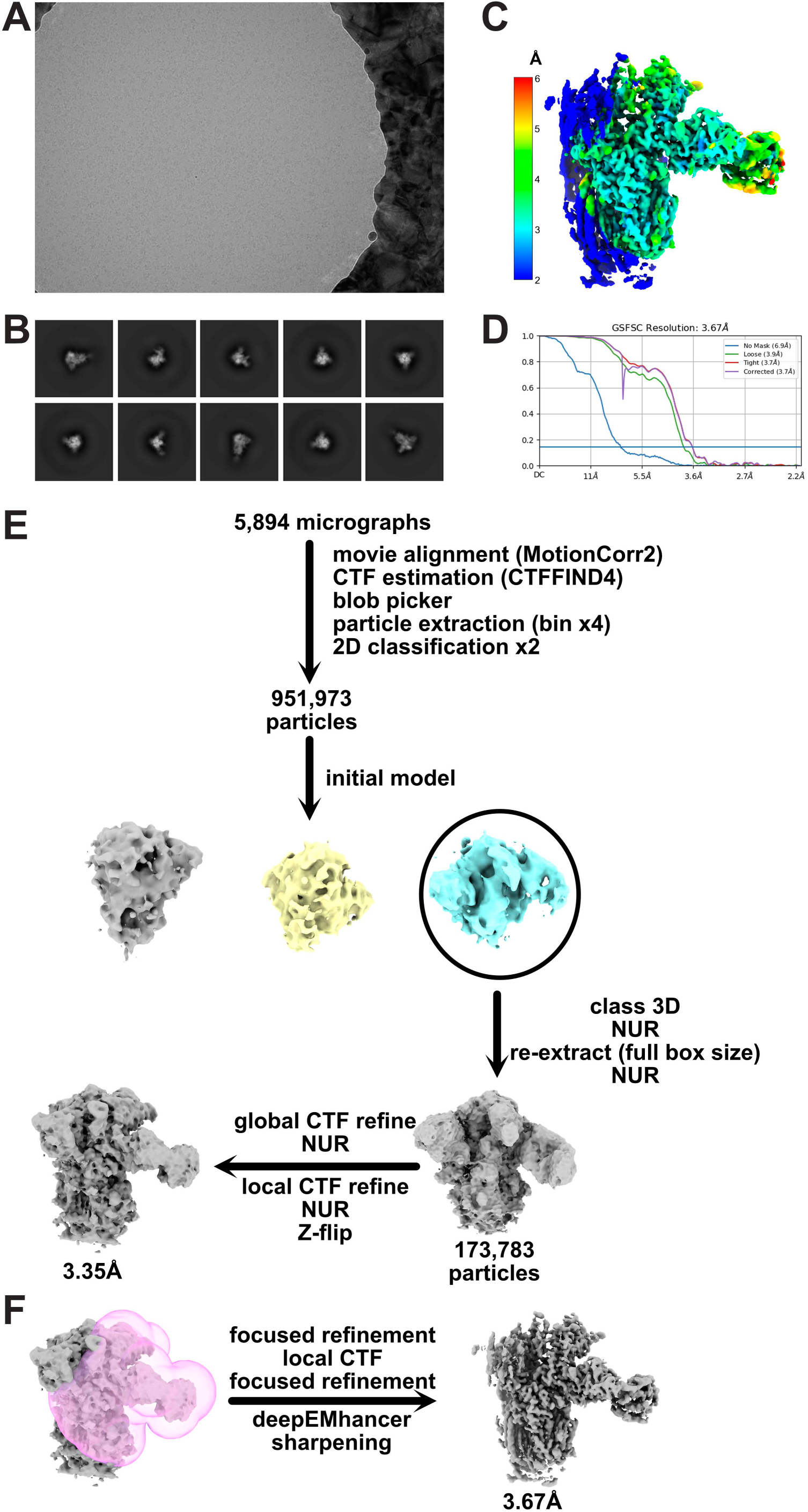
Cryo-EM data processing scheme and local resolution estimation of the final map. (**A**) Representative electron micrograph and (**B**) 2D class averages obtained for the SARS-CoV-2 Spike ectodomain in complex with PVI.V5-4 Fab. (**C**) Local resolution and (**D**) gold-standard Fourier shell correlation curves calculated with cryoSPARC v3.3.1. for the locally refined S1-Fab complex. Detailed cryo-EM data processing workflow for the global (**E**) and local (**F**) maps. See the Methods section for more details.

